# Polymorphisms in HIV-1 *nef* are associated with plasma concentration of biomarkers of endothelial activation

**DOI:** 10.1101/2022.10.10.511468

**Authors:** Genevieve Mezoh, Nereshni Lutchman, Eleanor M Cave, Katherine Prigge, Catherine Worsley, Neil Martinson, Elizabeth Mayne, Bronwen E Lambson, Penny L Moore, Nigel J Crowther

**Affiliations:** Department of Genome Sciences, School of Medicine, University of Washington, Seattle, Washington, United States of America; Department of Chemical Pathology, Faculty of Health Sciences, University of the Witwatersrand Johannesburg, South Africa; Department of Chemical Pathology, National Health Laboratory Service, Johannesburg, South Africa; Department of Immunology, University of Pretoria, Johannesburg, South Africa; Department of Molecular Medicine and Haematology, Faculty of Health Sciences, University of the Witwatersrand Johannesburg, South Africa; Perinatal HIV Research Unit (PHRU), University of the Witwatersrand Johannesburg, South Africa; Division of Immunology, Health Sciences Faculty, University of Cape Town, Cape Town, South Africa; Centre for HIV and STIs, National Institute for Communicable Diseases of the National Health Laboratory Service, Johannesburg, South Africa; SA MRC Antibody Immunity Research Unit, Department of Virology, Faculty of Health Sciences, University of the Witwatersrand Johannesburg, South Africa

**Keywords:** Endothelial activation, inflammation, HIV Nef, cardiovascular disease

## Abstract

Infection with HIV is associated with an increased risk of cardiovascular disease (CVD), which may be mediated by the effect of the viral proteins, Nef and Tat, on inflammation and endothelial activation. The viral genes coding for Nef and Tat contain numerous polymorphisms, which we hypothesised may be differentially associated with endothelial activation. Therefore, our aim was to assess the association of these polymorphisms with endothelial activation and inflammation in subjects infected with HIV-1.

The HIV-1 *nef* and *tat* genes were sequenced from clinical isolates from 31 and 34 patients, respectively. Plasma concentration of biomarkers of endothelial activation (intercellular adhesion molecule-1 (ICAM-1), vascular cell adhesion molecule-1 (VCAM-1), endothelial leukocyte adhesion molecule-1 (E-selectin), monocyte chemoattractant protein-1 (MCP-1) and von Willebrand factor (vWF)), and biomarkers of inflammation (tumor necrosis factor-α (TNF-α), interleukin-6 (IL-6) and interleukin-8 (IL-8)), were measured. Analysis of HIV-1 *nef* gene sequences identified five polymorphisms (V16I, H40Y, T50A,H, S169N and H188Q,S) that were each significantly (p<0.05) associated with ICAM-1 plasma concentration. An additive effect of these variants on plasma ICAM-1 concentration (p=0.004 for trend), was observed. No significant associations were seen between Tat amino acid residues and plasma concentration of markers of endothelial activation and inflammation. These are the first human *in vivo* data that support the hypothesis that *nef* gene polymorphisms impact endothelial function.

**Importance:** Cardiovascular disease (CVD) is a leading cause of mortality in adults living with HIV, which may in part be due to endothelial activation and inflammation caused by the viral proteins, Nef and Tat. However, there is no data from humans supporting the CVD-associated Nef and Tat hypothesis, and assays for accurately measuring Nef and Tat plasma concentrations are not currently available. Therefore, we hypothesized that polymorphisms in the *nef* and *tat* genes of clinical viral isolates may be associated with host plasma markers of endothelial activation and inflammation. Our results show that this was the case, with five *nef* polymorphisms showing both individual and additive association with plasma concentration of ICAM-1. The HIV-1 *Tat* gene, however, showed no significant association with plasma concentrations of markers of endothelial activation and inflammation. This is the first human study to directly link Nef to endothelial activation and to provide a possible screening tool i.e., *nef* genotyping, for identifying individuals at high risk of endothelial-based diseases.

## Introduction

Antiretroviral therapy (ART) has resulted in a marked decrease in AIDS-related deaths, as well as opportunistic infections, in people living with HIV (PLWH). However, non-communicable diseases such as cardiovascular diseases (CVDs) are now an emerging cause of concern with multiple studies showing an increased number of CVD-related deaths in PLWH compared to HIV-unaffected individuals (Chow et al., 2012, Freiberg et al., 2013). However, most of these reports stem from Europe and the US. The mechanism for this enhanced level of CVD in the context of HIV infection has not yet been defined and the prevalence of CVD among PLWH in sub-Saharan Africa has not been established. The direct role of ART as a cause of CVD is controversial with conflicting data (Obel et al., 2007, Lang et al., 2010, Arildsen et al., 2013), due to a gradual shift to less metabolically active ART regimens over time. Alternatively, HIV infection itself has been proposed to contribute towards the development of CVD through its effects on the vascular endothelium (Wang et al., 2015).

Endothelial dysfunction is regarded as a characteristic marker for the development of CVD (Münzel et al., 2008). The endothelium maintains the vasculature in a quiescent state by the production of nitric oxide (NO). This attenuates inflammation, thrombosis, and cell proliferation and migration. Damage to the endothelium, which may occur due to infection or turbulent blood flow leads to a decrease in NO production and activation of the endothelium (Liao, 2013). The activated endothelial cells the secrete a host of factors, including intercellular adhesion molecule-1 (ICAM-1), vascular cell adhesion molecule-1 (VCAM-1), endothelial leukocyte adhesion molecule-1 (E-selectin), monocyte chemoattractant protein-1 (MCP-1) and von Willebrand factor (vWF) (Verma et al., 2003, Flammer et al., 2012, Mezoh and Crowther, 2019) to restrict endothelial damage but in the presence of CVD risk factors such as hypercholesterolaemia and high blood pressure this response is augmented. Endothelial damage is also accompanied by an inflammatory response (Bruyndonckx et al., 2013), which involves the upregulation of cytokines such as tumor necrosis factor-α (TNF-α), interleukin-6 (IL-6) and interleukin-8 (IL-8) (De Pablo-Bernal et al., 2014). Chronic activation of the endothelium results in a dysfunctional state, which can lead to vascular damage, including the progression of atherosclerosis (Deanfield et al., 2007). Studies conducted on PLWH show that endothelial activation, as assessed using plasma biomarkers such as VCAM-1 and ICAM-1, demonstrate a broad concentration of these proteins that overlap with those in HIV-unaffected subjects (Graham et al., 2013, Fourie et al., 2015, Mezoh et al., 2021). This differential effect of the virus on endothelial activation may arise from either the genetic heterogeneity of the virus or heterogeneity of the host response to the infection.

The HIV genome exhibits a very high mutation rate and substantial diversity (Sanjuán et al., 2010). Variation in HIV viral protein sequences has been shown to influence the host response to the virus. As an example, *tat* sequence variation has been associated with variation in chemoattractant activity and ability to stimulate TNF-α production (Siddappa et al., 2006, Campbell et al., 2007). Cohort studies have also implicated sequence variation with clinical outcomes. Genetic variation in *tat* has been associated with neurological disorders in people living with HIV (Mayne et al., 1998, Cowley et al., 2011, Li et al., 2012). Similarly, sequence analysis of HIV-1 *tat* isolated from PLWH with and without dementia revealed an association between signature residues and neurological impairment (Bratanich et al., 1998). In addition, sequence analysis of the HIV-1 *nef* gene isolated from a cohort of HIV-affected, ART-naïve infants and children with varying stages of disease progression, revealed unique polymorphisms specific to slow progressors at the N-terminal and C-terminal domains (Walker et al., 2007).

Several studies support the hypothesis that HIV viral proteins may be role players in the development of CVD via their effects on the vascular endothelium (Dhawan et al., 1997, Matzen et al., 2004, Wang et al., 2014). Thus, *in vitro* and *in vivo* studies have shown that the HIV viral proteins, Nef (Wang et al., 2014), Tat (Dhawan et al., 1997, Matzen et al., 2004) and gp120 (Jiang et al., 2010), upregulate the expression of markers of endothelial activation and inflammation. The HIV-1 Nef and Tat amino acid sequences have been extensively studied in relation to HIV disease progression, however, limited research has been performed on these proteins within the context of non-communicable diseases. Almodovar *et al*. revealed

HIV-1 Nef polymorphisms to be associated with pulmonary hypertension (Almodovar et al., 2012). Based on these observations, we hypothesized that polymorphisms in HIV-1 Nef and Tat may be associated with endothelial activation in PLWH. Therefore, the aim of this study was to sequence *nef* and *tat* genes from clinical HIV-1 isolates, and determine whether mutations in these viral genes were associated with plasma concentration of markers of endothelial activation (VCAM-1, ICAM-1, E-selectin, vWF, MCP-1) and inflammation (IL-6, TNF-α, IL-8).

## Materials and Method

### HIV-1 clinical isolates

Eighty (80) PLWH, ART-naïve Black South African individuals between the ages of 30 to 50 years were recruited from the Nthabiseng and Zazi Clinics, Chris Hani Baragwanath Hospital, as described in our previous study (Mezoh et al., 2021). Subjects with clinical conditions known to influence endothelial function, as defined in Mezoh et al., 2021, were excluded from this study. Subjects were screened for HIV at a walk-in point of care HIV testing service, using the Bioline HIV 1/2 3.0 rapid test kit (Standard Diagnostics Inc, Gyeonggi-do, Republic of Korea) (Govender et al., 2014) and HIV seropositivity was confirmed using the Cobas® HIV-1 quantitative nucleic acid test on the Cobas® automated 6800 system. Plasma samples were obtained from blood collected in EDTA tubes and stored at −80°C until further analysis. These samples were used for the measurement of markers of endothelial activation (ICAM-1, VCAM-1, E-selectin, MCP-1 and vWF) and inflammation (TNF-α, IL-6 and IL-8).

### Clinical and blood analyte measurements

Blood pressure measurements, height, weight and body mass index (BMI) were measured using previously described methods (Crowther and Norris, 2012). Viral loads were measured from EDTA plasma samples using the Cobas® HIV-1 quantitative nucleic acid test on the Cobas® automated 6800 system, while CD4 count was measured from EDTA plasma samples by flow cytometry, following standard protocols (Glencross et al., 2008). Plasma concentration of ICAM-1, VCAM-1, E-selectin, MCP-1, TNF-α, IL6 and IL8 were measured using a customised Human Magnetic Luminex Screening Assay kit (R&D Systems, Minneapolis, MN, U.S.A) on the Bio-Plex®Multiplex System (Bio-Rad, Hercules, CA, U.S.A) with Luminex xMAP technology, according to the manufacturer’s instructions. The vWF was measured by ELISA using a human vWF ELISA kit (Merck, Darmstadt, Germany) as described by the manufacturer.

The study was approved by the University of the Witwatersrand’s Human Research Ethics Committee (Medical) and all participants provided written informed consent prior to their inclusion in this study.

### Sequencing of HIV-1 *nef* and *tat* viral genes

Viral RNA from EDTA plasma samples was extracted using a QIAamp viral RNA minikit (Qiagen, Hilden, Germany) according to the manufacturer’s instructions. Synthesis of cDNA was carried out using SuperScript III reverse transcriptase (Invitrogen, Waltham, MA, U.S.A.) and the QuantiTect Reverse Transcription kit (Qiagen) with primers Vif1: 5’-GGGTTTATTACAGGGACAGCAGAG-3’ and OFM19: 5’-GGTAGGATCTCTACAATACTTGGCACTG-3’ according to the manufacturer’s protocol. The *nef* and *tat* genes were amplified in a nested polymerase chain reaction (PCR) using the high fidelity Platinum^TM^ *Taq* DNA polymerase (Invitrogen, CA, U.S.A) and HIV-1 subtype C *nef* and *tat* specific outer primers as described by Salazar-Gonzalez *et al*. (Salazar-Gonzalez et al., 2011), and inner primers as described by Bredell *et al*., (Bredell et al., 2007) with slight modifications. Outer primers Vif1 and OFM19 were used to amplify the *tat* exon 1 and *tat* exon 2 fragments in the first PCR reaction, which amplified the full *env* encoding region. In the second PCR reaction, *tat* exon 1 was amplified using inner primers Tat1F2: 5’-GGTAGGATCTCTACAATACTTGGCACTG-3’ and N1156R2: 5’-TCATTGCCACTGTCTTCT GCTCT-3’ generating a 216bp product, while *tat* exon 2 was amplified using inner primers Nf: 5’-TGACCTGGATGCAGTGG-3’ and CT1re: 5’-GACTTCCCAGATACTTAAGAGCTTCCCACC-3’ yielding an 87bp amplicon. The 639bp *nef* fragment was amplified using outer primers NefOF: 5’-GTGGAATTCCTGGGACG-3’ and NefOR: 5’-AGGCAAGCTTTATTGAGG-3’ in the first PCR reaction, and inner primers NefIF: 5’-CCTAGAAGAATAAGACAGGGC-3’ and NefIR: 5’-CTTATATGCAGCAT CTGAGG-3’ in the second PCR reaction. The *nef* and *tat* encoding regions were sequenced using the inner primer sets with the ABI PRISM Big Dye Terminator Cycle Sequencing Ready Reaction kit (Applied Biosystems, Waltham, MA, U.S.A.) according to the manufacturer’s instruction and resolved on an ABI 3100 automated genetic analyzer. Sequencing reactions of the HIV-1 *nef* and *tat genes* from each individual of the PLWH cohort were performed in duplicate.

### Bioinformatic analysis of sequencing data

The nucleotide sequence regions coding for the HIV-1 Nef and Tat proteins were identified and assembled using the CLC workbench (CLC Bio, Aarhus, Denmark) and edited with BioEdit (ver. 5.0.9; Informer Technologies Inc., Los Angeles, CA, U.S.A.). The quality of sequencing data was assessed using FastQC (Babraham Bioinformatics, Cambridge, U.K.). Nucleotide sequences were translated to their respective amino acid sequences, following which, multiple sequence alignments were performed using BioEdit. HIV-1 n*ef* and *tat* amino acid sequences were analysed and compared against sequence sets retrieved from the Los Alamos HIV-1 Sequence Database. Phylogenetic trees were constructed using MEGA to assess linkages and eliminate contaminations. The HIV-1 Nef and Tat functional domains were annotated using the HIV-1 proteomics resource (Doherty et al., 2005). The HIV-1 Nef amino acid residues were numbered following HIV-1 HXB2 Nef which is the general sequence used for numbering.

The HIV-1 consensus C Nef differs in number from the HXB2 Nef by the addition of an extra E amino acid residue at position 62, resulting in a shift in numbering by 1 residue from position 61.

### Statistical analysis

Data that was not normally distributed was expressed as median [interquartile range] while data with a normal distribution was expressed as mean ± SD. The HIV-1 amino acid sequences were aligned to the HXB2 reference strain obtained from the Los Alamos HIV-1 Sequence Database to identify Nef and Tat variants. For each variant, subjects were grouped according to the amino acid sequence and plasma concentration of VCAM-1, ICAM-1, MCP-1, E-selectin, vWF, IL-6, IL-8 and TNF-α were compared between the groups using either a Mann-Whitney U test or Kruskal-Wallis ANOVA, depending on the number of sequence variants found at each polymorphic site. Statistical analysis was performed using GraphPad prism 7 (GraphPad Software Inc., San Diego, CA, U.S.A.) and Statistica software v13 (Statsoft Inc., Tulsa, OK, U.S.A.).

## Results

### Clinical and virologic characteristics of the study group

Out of the 80 recruited PLWH ARV-naïve participants, sufficient plasma sample volume for sequencing was available for 65 (81.3%) subjects. Viral RNA was extracted from all these samples, and HIV-1 *nef* and *tat* sequences were successfully amplified from viral RNA extracted from 30/65 and 34/65 patients, respectively. The mean age of this sub-group of 35 subjects was 39 years, with a mean CD4 T cell count of 376 cells/mm^3^, mean viral load of 116415 copies/mL and a mean BMI of 26.4 Kg/m^2^. The low success rate for *nef* and *tat* amplification is likely due to the fact that 17 out of the 65 PLWH adults had a viral load level below 1,500 cells/mm^3^.

### Comparison of cardiometabolic variables between groups

We compared data for the subjects from whom viral DNA sequences were obtained, with those from whom no sequence data was obtained to assess whether the sequenced group appeared to differ from the latter group. No significant differences were observed between these groups for age, BMI, blood pressure, TNF-α, IL-6, ICAM-1, E-selectin, vWF and MCP-1 concentration (Table 1). However, more males, higher concentration of viral load and VCAM-1, and lower concentration of CD4 counts and IL-8 were observed in the sequenced compared to the non-sequenced group.

**Table 1:** Comparison of HIV sequenced and non-sequenced groups.

| Variables | HIV non-sequenced (n=45) | HIV sequenced (n=35) |
| --- | --- | --- |
| **Age (years)** | 36.71 ± 6.93 | 38.46 ± 13.31 |
| **Gender, male n (%)** | 5 (11.11) | 11 (31.42)* |
| **BMI (kg/m<sup>2</sup>)** | 26.86 ± 4.64 | 26.41 ± 9.38 |
| **Waist circumference (cm)** | 88.89 ± 9.66 | 84.46 ± 13.31 |
| **Diastolic BP (mmHg)** | 80.33 ± 15.22 | 79.71 ± 13.46 |
| **Systolic BP (mmHg)** | 137.07 ± 20.15 | 129.51 ± 21.93 |
| **Glucose (mmol/L)** | 4.66 ± 1.71 | 4.38 ± 0.35 |
| **Triglyceride (mmol/L)** | 0.87 [0.61, 1.10] | 0.58 [0.38, 0.76]*** |
| **Total cholesterol (mmol/L)** | 3.89 ± 0.84 | 3.22 ± 0.78** |
| **LDL-C (mmol/L)** | 2.31 ± 0.62 | 1.93 ± 0.66* |
| **HDL-C (mmol/L)** | 1.08 [0.91, 1.35] | 0.92 [0.71, 1.20] |
| **CD4+ cell count (cells/mm<sup>3</sup>)** | 542.56 ± 314.68 | 375.69 ± 241.17* |
| **Viral load (copies/mL)** | 8981.22 ± 18465.38 | 116414.57 ± 94695.18*** |
| **TNF-α (pg/mL)** | 6.2 [4.5, 9.6] | 7.7 [5.2, 10.5] |
| **IL-6 (pg/mL)** | 2.9 [1.3, 4.6] | 3.9 [2.3, 5.3] |
| **IL-8 (pg/mL)** | 8.4 [5.2, 24.1] | 6.0 [3.2, 7.9]** |
| **ICAM-1 (ng/mL)** | 464.8 [344.1, 923.1] | 848.4 [403.4, 1048.7] |
| **VCAM-1 (ng/mL)** | 522.9 [426.3, 805.6] | 954.3 [669.8, 1477.7]* |
| **E-selectin (ng/mL)** | 27.9 [19.8, 33.9] | 30.0 [23.4, 44.1] |
| **vWF (µg/mL)** | 20.7 [12.1, 29.9] | 26.9 [18.4, 43.9] |

| **MCP-1 (pg/mL)** |  |  | 139.3<br>[114.9,<br>181.1] | 137.2<br>[108.5,<br>209.7] |
| --- | --- | --- | --- | --- |
| Variables | HIV non-sequenced<br>(n=45) | HIV sequenced<br>(n=35) |  |  |
| **Age (years)** | 36.71 ± 6.93 | 38.46 ± 13.31 |  |  |
| **Gender, male n (%)** | 5 (11.11) | 11 (31.42)* |  |  |
| **BMI (kg/m<sup>2</sup>)** | 26.86 ± 4.64 | 26.41 ± 9.38 |  |  |
| **Waist circumference (cm)** | 88.89 ± 9.66 | 84.46 ± 13.31 |  |  |
| **Diastolic BP (mmHg)** | 80.33 ± 15.22 | 79.71 ± 13.46 |  |  |
| **Systolic BP (mmHg)** | 137.07 ± 20.15 | 129.51 ± 21.93 |  |  |
| **Glucose (mmol/L)** | 4.66 ± 1.71 | 4.38 ± 0.35 |  |  |
| **Triglyceride (mmol/L)** | 0.87 [0.61, 1.10] | 0.58 [0.38, 0.76]*** |  |  |
| **Total cholesterol (mmol/L)** | 3.89 ± 0.84 | 3.22 ± 0.78** |  |  |
| **LDL-C (mmol/L)** | 2.31 ± 0.62 | 1.93 ± 0.66* |  |  |
| **HDL-C (mmol/L)** | 1.08 [0.91, 1.35] | 0.92 [0.71, 1.20] |  |  |
| **CD4+ cell count (cells/mm<sup>3</sup>)** | 542.56 ± 314.68 | 375.69 ± 241.17* |  |  |
| **Viral load (copies/mL)** | 8981.22 ± 18465.38 | 116414.57 ± 94695.18*** |  |  |
| **TNF-α (pg/mL)** | 6.2 [4.5, 9.6] | 7.7 [5.2, 10.5] |  |  |
| **IL-6 (pg/mL)** | 2.9 [1.3, 4.6] | 3.9 [2.3, 5.3] |  |  |
| **IL-8 (pg/mL)** | 8.4 [5.2, 24.1] | 6.0 [3.2, 7.9]** |  |  |
| **ICAM-1 (ng/mL)** | 464.8 [344.1, 923.1] | 848.4 [403.4, 1048.7] |  |  |
| **VCAM-1 (ng/mL)** | 522.9 [426.3, 805.6] | 954.3 [669.8, 1477.7]* |  |  |
| **E-selectin (ng/mL)** | 27.9 [19.8, 33.9] | 30.0 [23.4, 44.1] |  |  |
| **vWF (µg/mL)** | 20.7 [12.1, 29.9] | 26.9 [18.4, 43.9] |  |  |
| **MCP-1 (pg/mL)** | 139.3 [114.9, 181.1] | 137.2 [108.5, 209.7] |  |  |

Data given as mean ± standard deviation or median [interquartile range]; n indicates the number of samples; \*p<0.05, \*\*p<0.005, \*\*\*p<0.0005 vs HIV non-sequenced.

### Sequence analysis of HIV-1 Nef and Tat

An HIV-1 Nef sequence alignment of the 30 clinical isolates is shown in Figure 1. The N-terminal and C-terminal domains spanning position 1 to 79 and 165 to 220 respectively, shows substantial variability in the HIV-1 Nef consensus C sequence, with the centre region from position 80 to 164 remaining relatively conserved. In contrast to Nef, the HIV-1 Tat multiple sequence alignment of the clinical isolates from the HIV-positive subset (Figure 2) showed that variations in the HIV-1 Tat consensus C sequence was largely located at the C-terminal domain, with the N-terminal domain and central region remaining relatively conserved.

**Figure 1:**
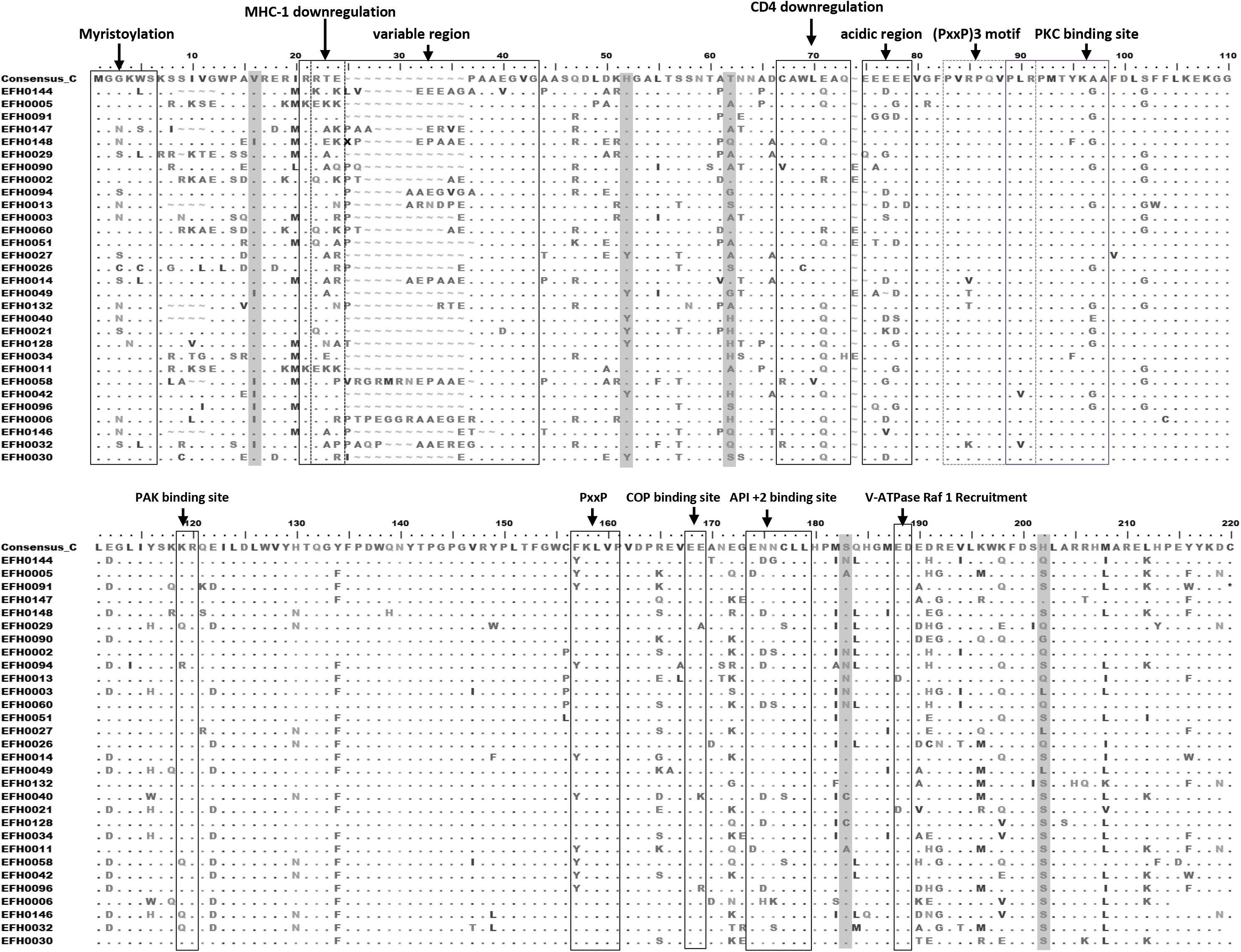
Multiple sequence alignment of plasma derived HIV-1 Nef protein sequences. Nucleotide sequences were aligned with the HIV-1 *nef* Clade C consensus sequence (retrieved from Los Alamos National Laboratory HIV Sequence Database) using BioEdit, translated and numbered following HXB2. Sequences have been placed in decreasing order of ICAM-1 concentration. Functional domains, which include the sequence regions involved in Nef trafficking and internalization resulting in MHC-1 and CD4 downregulation and other functional motifs, such as those involved in Nef modification and signalling, which includes the region for myristoylation, the (PxxP)3 motif, the PKC, PAK, COP and API +2 binding sites, are all boxed. Amino acid residues determined to be associated with ICAM-1 concentration is shaded in grey. Other annotated indels are boxed. Amino acid residues that are the same as that of Nef Clade C are represented in dots (.), while gaps in the alignment are represented with tildes (~).

**Figure 2:**
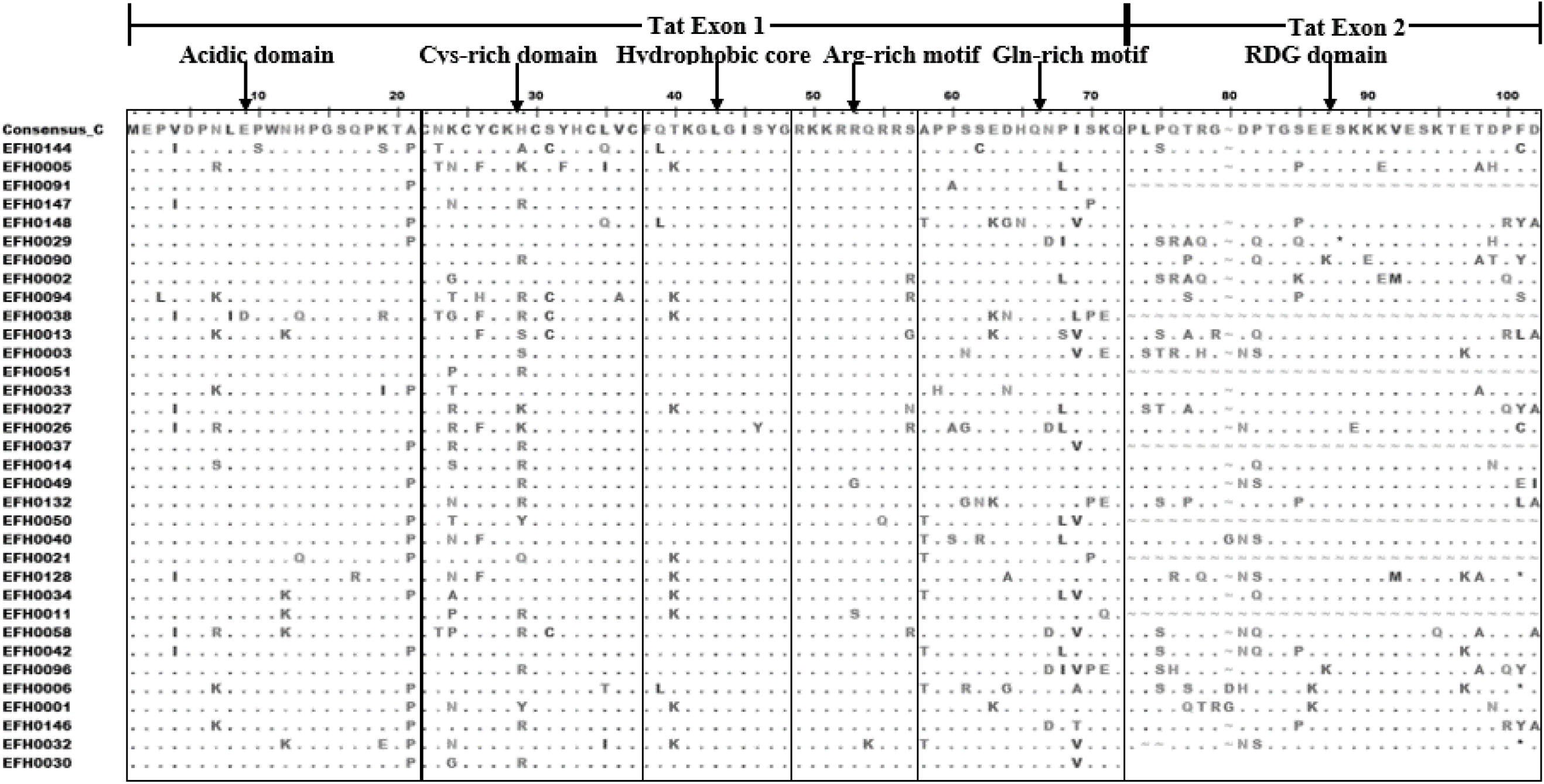
Multiple sequence alignment of plasma derived HIV-1 Tat sequences. Sequences were aligned with the HIV-1 *tat* Clade C consensus sequence (retrieved from Los Alamos) using BioEdit, translated and numbered following HXB2. Sequences have been placed in decreasing order of ICAM-1 concentration. The functional domains which include the acidic region, cysteine-rich motif, hydrophobic core region, arginine-rich motif, glutamine-rich motif and RDG domain, are all boxed (Mahlknecht et al., 2008). Amino acid residues that are the same as that of HIV-1 Nef Clade C are represented in dots (.) while gaps in the alignment are represented with tildes (~).

### Association of Nef and Tat variants with plasma biomarker concentration of endothelial activation and inflammation

Sequence analysis of amino acid residues of HIV-1 Nef (Figure 1) revealed several statistically significant associations between sequence variants and markers of endothelial activation and inflammation. The consensus amino acids 16V, 40H and 50T, were significantly associated with higher concentrations of ICAM-1, while mutations at codon 169 (from Ser to Asn) and 188 (from His to Gln), were also associated with higher concentrations of ICAM-1 (Figure 3 A-E). With regard to this latter association, the HIV-1 Nef consensus C amino acid residue at codon 188 (His) was present in only one subject out of the 30 sequenced samples with 20% (6/30) of the study population having the variant Gln, and 63% (19/30) having the variant Ser. In addition, the consensus amino acids 182K and 205D were associated with higher concentrations of MCP-1 and TNF-α, respectively and a mutation at codon 202 from Tyr to Phe was associated with increased concentration of VCAM-1 (Table 2).

**Figure 3:**
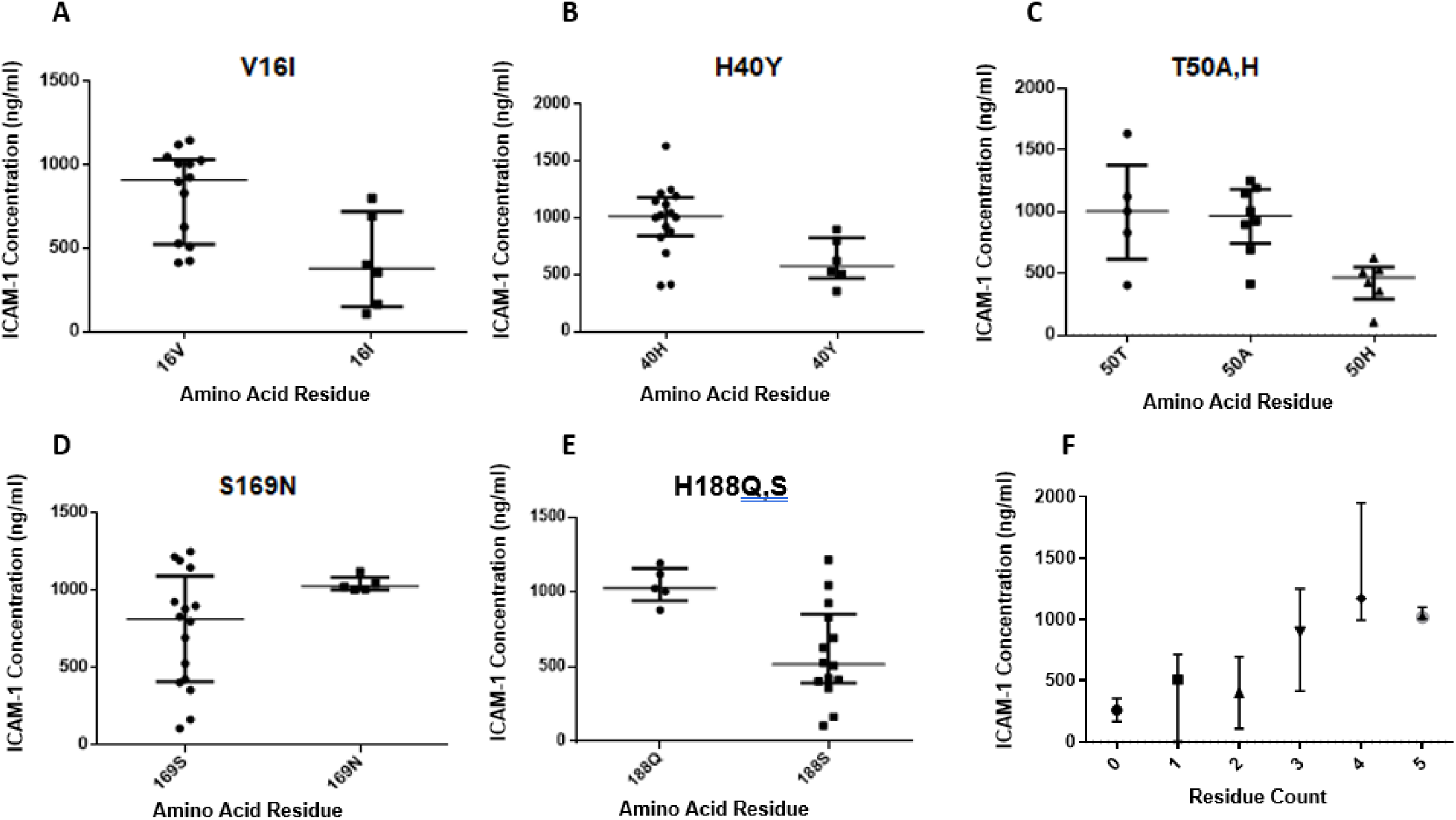
HIV-1 Nef amino acid residues associated with increased concentration of plasma ICAM-1. (A-E) Dot plot of ICAM-1 concentration versus amino acid residue at each of the 5 polymorphic loci that were associated with ICAM-1. Data expressed as median with interquartile range; graphs A to E all show data that are significantly different between groups at *p* < 0.05. The number of Nef ICAM-1-associated amino acid residues in each patient was counted (residue count) and subjects grouped according to the number of ICAM-1-associated residues and plotted against their ICAM-1 concentrations; p=0.004 from ANOVA (F).

**Table 2:** HIV-1 Nef polymorphisms significantly associated with plasma concentration of VCAM-1, MCP-1 and TNF-α.

| **Biomarker** | **Variant** | **Amino acid** | **N** | **Biomarker level (median [interquartile range] (ng/mL))** | **p value** |
| --- | --- | --- | --- | --- | --- |
| **VCAM-1** | Y202F | Y | 17 | 847 [506, 1324.55] | **0.020** |
|  |  | F | 9 | 1293 [1173, 2337] |  |
| **MCP-1** | K182M | K | 18 | 150 [115, 217] | **0.048** |
|  |  | M | 8 | 103 [93, 130] |  |
| **TNF-α** | D205N | D | 25 | 8.27 [6.70, 11.0] | **0.037** |
|  |  | N | 5 | 5.21 [4.06, 6.64] |  |
|  |  | K | 5 | 1478 [1203, 1617] |  |

Data given as median [lower quartile, upper quartile]; N indicates the number of individuals

Sequence analysis of HIV-1 Tat showed no association of any of the variants with plasma concentration of markers of inflammation or endothelial activation.

To determine if there was an additive effect of these variants on ICAM-1 concentration, the number of ICAM-1-associated Nef amino acid residues (residue count, range of 0 to 5; N=2,9,2,7,6 and 4 respectively) present in the viral isolates of each patient were counted and plotted against the plasma concentration of ICAM-1 (Figure 3F). The level of ICAM-1 increased with the number of ICAM-1-associated Nef amino acid residues (p=0.004 for trend by ANOVA).

## Discussion

In this study we investigated whether HIV infection was linked to endothelial activation via the action of the viral proteins Nef and Tat. This was achieved by sequencing the *nef* and *tat* genes from clinical isolates following which the amino acid residues at the polymorphic loci were screened for association with plasma concentration of markers of endothelial activation and inflammation. A total of five variants were identified in the Nef protein that were linked to increased concentration of ICAM-1. In addition, our analysis showed an additive effect of these variants on ICAM-1 concentration. Three Nef variants were also observed to be each associated with plasma concentrations of VCAM-1, TNF-α and MCP-1. No HIV-1 Tat sequence variants were linked to increased concentration of any of the biomarkers measured.

We found that the HIV-1 Nef amino acid residues 16V, 40H and 50T at the N-terminal domain, and 169N and 188Q at the C-terminal domain were linked with increased concentration of ICAM-1, whilst the Nef residues 202F, 182K and 205D were linked with higher concentration of VCAM-1, MCP-1 and TNF-α, respectively. A study by Walker et al (2007) reported that mutations in HIV-1 *nef* were linked to a loss in MHC-1 downregulation and other functions involving cell signaling and these variants were also present at the N- and C-terminal domains of the protein. The N-terminal domain of HIV-1 Nef contains regions involved in MHC-1 and CD4 downregulation, while the C-terminal domain contains regions involved in the internalization of the Nef protein (Mandic et al., 2001). *In vitro* studies conducted by Mandic *et al*. (2001) using simian immunodeficiency virus (SIV) showed the amino acid residues 39Y, 194L, 195M, 204D and 205D to be a prerequisite for the cellular uptake of Nef by Jurkat and 293-T cells. Changes to any of the consensus amino acid residues resulted in a loss in infectivity. A substitution to the amino acid residue 205D has thus been linked to a decreased uptake of the Nef protein by cells (Mandic et al., 2001), and slow disease progression (Walker et al., 2007). In the current study, the D205N mutation was found to be associated with lower plasma concentration of TNF-α, and based on the study by Mandic et al. (2001), this may be related to reduced Nef uptake by host immune cells.

In our study, multiple Nef variants were associated with ICAM-1 plasma concentration. This is of interest because Nef is reported to play a direct role in the upregulation of ICAM-1 via the extracellular signal-regulated kinase (ERK) mitogen-activated protein kinase (MAPK) signalling pathway (Fan et al., 2010). Phosphorylation of ERK1 and ERK2 is enhanced by Nef, which activates the ERK/MAPK pathway resulting in the expression of the adhesion molecule ICAM-1. This is demonstrated by *in vitro* studies conducted by Fan et al. (2010), in which the HIV-1 Nef-induced overexpression of ICAM-1 by human vascular endothelial cells was inhibited by the kinase inhibitor, PD98059, a chemical known to inhibit the ERK/MAPK pathway.

Studies have also shown associations between Nef and CVD in HIV-positive subjects (Wang et al., 2014, Stolp et al., 2012, Almodovar et al., 2011). In a study conducted by Almodovar et al. (2012), HIV-1 Nef sequences from 11 PLWH with pulmonary hypertension were compared to sequences obtained from 7 PLWH without pulmonary hypertension, and polymorphisms in Nef were found to be associated with the presence of this disease (Almodovar et al., 2012). The Nef mutations identified in the PLWH with pulmonary hypertension were R19K, R21Q, H40Y, A53P, L58V, E63G, M79I, T80N, Y81F and P150A,R,S,K,Q, (Almodovar et al., 2012). However, in a validation cohort from the U.S.A consisting of 11 PLWH with pulmonary hypertension compared against 22 PLWH without pulmonary hypertension, only R21Q, H40Y, A53P, L58V, E63G, Y81F and P150A,R,S,K,Q, mutations were validated (Almodovar et al., 2012). Given that 40H was associated with pulmonary hypertension, and was also associated with ICAM-1 concentration in the current study, this variant could be a key player in the aetiology of CVD in PLWH. The current study also showed that subjects carrying viruses with higher numbers of the five ICAM-1-associated residues had higher plasma concentrations of ICAM-1. This suggests an additive effect of the variants on ICAM-1 concentration.

A limitation of this study is the low number of HIV-1 Nef and Tat amino acid sequences that were successfully sequenced. In this study population, 26 % had viral load levels below the threshold for successful sequencing (Dudley et al., 2012). Several other factors could possibly account for the low sequencing success rate. This includes poor sample quality, degraded viral RNA, insertions/deletions within the HIV genome as well as the potential of obtaining a mix of PCR products thus hindering sequencing due to insertions and deletion as not all fragments would be of the same length, A similar observation was reported by Rousseau et al (2006) in which near full-length genomes of HIV-1 subtype C was successfully sequenced from 67 % of samples. A sample size calculation was not performed for this study because *nef* and *tat* are very polymorphic genes and we had no prior information on which of the variants could be associated with the selected plasma analytes. Another limitation is the discrepancy between the HIV-sequenced and unsequenced population. The sequenced population had a higher viral load and concentration of VCAM-1, and lower CD4 cell counts and IL-8 concentration. As these results were obtained from a small dataset, it is difficult to determine whether they are representative of all PLWH. This warrants expansion to a larger cohort of PLWH. In addition, functional studies need to be performed to determine how the identified *nef* polymorphisms lead to alterations in ICAM-1 concentration.

The increasing levels of non-communicable diseases, such as CVD, in the HIV affected population warrants further investigation to identify interventions that could attenuate this high CVD risk. To the best of our knowledge, this is the first study to investigate HIV-1 Nef and Tat amino acid sequences from clinical isolates in the context of endothelial activation and its aetiological role in CVD. We found amino acid residues in HIV-1 Nef that were associated with endothelial activation. Despite the ability of anti-retroviral therapies to decrease viral load in PLWH, the HIV Nef protein is seen to persist in individuals with undetectable viral load (Raymond et al., 2019), and it is therefore possible that it may continue to exert its action on endothelial cells even in subjects who are virally suppressed. Therefore, it is possible that blocking Nef activity, specifically in subjects carrying the Nef variants identified in this study, could be a potential therapeutic strategy to alleviate endothelial activation and associated vascular disease. Future work in this field would include the development of Nef inhibitors, possibly anti-Nef antibodies, and using such agents in *in vitro* and *in vivo* HIV-1 Nef inhibitory studies, to assess if inhibiting this viral protein results in decreased endothelial cell activation. It would also be important to determine whether Nef variants are associated with disease end points such as coronary artery disease, peripheral artery disease and stroke, all of which are associated with endothelial dysfunction.

## Acknowledgments

Funding to support this project was obtained from the South African National Health Laboratory Service (NHLS), the South African National Research Foundation (NRF), the Poliomyelitis Research Foundation (PRF) and the University of the Witwatersrand. The protocol was conceived by NJC, PLM and GM, while experiments were carried out by GM, who also analysed the data and wrote the paper under the supervision of PLM and NJC. The final paper was read and approved by all contributing authors. P.L.M. is supported by the South African Research Chairs Initiative of the Department of Science and Technology and the NRF (Grant No 98341). We acknowledge funding received from the Organization for Women in Science for the Developing World (OWSD) and Swedish International Development Cooperation Agency (SIDA) under the OWSD postgraduate training fellowship awarded to GM, as well as funding received from the Schlumberger Foundation and American Association for the Advancement of University Women postdoctoral fellowship awarded to GM.

